# Spexin modulates molecular thermogenic profile of adipose tissue and thermoregulatory behaviors

**DOI:** 10.1101/2021.09.10.456868

**Authors:** Shermel B. Sherman, Niraj Gupta, Mitchell Harberson, Riley Powers, Rebecca Rashleigh, Ramya Talla, Ashima Thusu, Jennifer W. Hill

## Abstract

Thermoregulation is a physiological process by which a mammal regulates body temperature in response to its environment. Within the human body, thermoregulatory behaviors and metabolism are modulated by circulating metabolic factors. In our study, we tested the ability of the neuropeptide spexin, which shares sequence homology to galanin, to regulate these functions in female mice. Supraphysiological levels of spexin in C57BL/6 mice were insufficient to protect against diet-induced obesity after 50 days of treatment. Behavioral analysis of long-term spexin treatment appeared to modulate anxiety-like behaviors by promoting exploratory behaviors and thermoregulatory behaviors of nest building that ceased when animals were housed at thermoneutral temperatures. Upon examination of the molecular profile of brown and white adipose tissue, treatment disrupted the thermogenic profile of white adipose tissue, in which β3-adrenergic receptor expression was downregulated. Our results reveal novel functions for spexin as a modulator of thermoregulatory behaviors and adipose tissue metabolism.

**Highlights:**

- Spexin treatment did not protect against diet-induced obesity in female mice.
- Spexin-treatment promoted thermoregulatory behaviors of nest building.
- Behaviors normalized when animals were housed in thermoneutral temperatures.

**Funding Sources:** Not applicable

**Disclosure Summary:** Nothing to disclose

## INTRODUCTION

Thermoregulation is the process by which a mammal regulates body temperature in response to the environment. It consists of both physiological responses and behavioral adaptations. For instance, when exposed to cold stress, animals respond by engaging in thermoregulatory behaviors such as nest building for warmth [1]. Similarly, heat generation is activated during hibernation allowing sleeping mammals to maintain their core body temperature [2]. In addition, body temperature can be manipulated to reduce energy use; for example, core body temperature drops with caloric restriction to reduce weight loss [3, 4].

The brain coordinates responses to thermal stress via sympathetic innervation and agonism of adrenergic receptors in two adipose depots: brown and white adipose tissue. In newborns, brown adipose tissue (BAT) is important for shivering thermogenesis and is found in surrounding the neck, interscapular region, and axillary [5–7]. In humans, BAT deposition is less extensive than in newborns but may contribute to weight loss [8–11]. Adrenergic receptors localized on the cell surface of brown adipocytes [12–17, 11, 18] are activated by norepinephrine released from sympathetic nerve terminals in adipose tissue. When adrenergic receptors are activated in BAT, uncoupling protein 1 (UCP1) synthesis and expression increases in mitochondria, which increases the transport of hydrogens into the mitochondrial matrix [19–22, 8, 9, 23, 24]. Besides increasing heat production, this process improves the insulin sensitivity of the tissue [25, 20, 8, 10]. The activity in BAT triggers lipolysis in white adipose tissue depots to release free-fatty acids into circulation that is used as energy within the BAT [19, 8, 26, 10]. The loss of adrenergic receptors can therefore disrupt thermogenesis and the potential for weight loss [27, 28].

Homeostatic mechanisms in the brain trigger physiological and behavioral changes utilized by an organism to maintain body temperature. Galanin, a neuropeptide of 30 amino acids, is involved in both metabolic processes of thermoregulation and nest building [29]. Within the last decade, spexin (SPX) a peptide hormone with less than 50% sequence homology to galanin was identified as a potential therapeutic target for the treatment of obesity [30, 31]. It also functions as an anxiolytic in male animals across different species [32–34]. Spexin binds to galanin receptor 2 (Galr2) and galanin receptor 3 (Galr3), which are expressed in the brain [35–37]. Spexin immunostaining in normal rat tissues revealed that spexin is also expressed throughout the CNS with intense immunostaining detected within the hypothalamus, cerebellum, striatum, and hippocampus [38]. Treatment with a modified spexin agonist for Galr2 revealed neuronal activation in the DRN, VTA, SN, dentate gyrus, central amygdala, prefrontal cortex, and nucleus accumbens [39].

Given this evidence, we hypothesized that spexin may function as a modulator of both adipogenesis and behavior. Using a female model of hyperspexinemia, we found that spexin disrupts the thermogenic profile of adipose tissue and promotes nest building as a thermoregulatory behavior.

## RESULTS

### Hyperspexinemia does not protect against diet-induced obesity in young adult female mice

Previous spexin studies in rodents revealed that between 30-50 days of spexin treatment with 25 μg/kg spexin was sufficient to protect rodents against diet-induced obesity [40, 30]. In our study, we challenged female C57BL/6 mice with 60% kcal fat high-fat diet for 50 days and administered 25 μg/kg spexin once daily by subcutaneous bolus injection (Figure 1a). Although previous studies revealed significant weight loss in spexin-treated animals, these supraphysiological levels of spexin were unable to protect female mice against diet-induced obesity (Fig. 1b, c).

**Fig 1.**
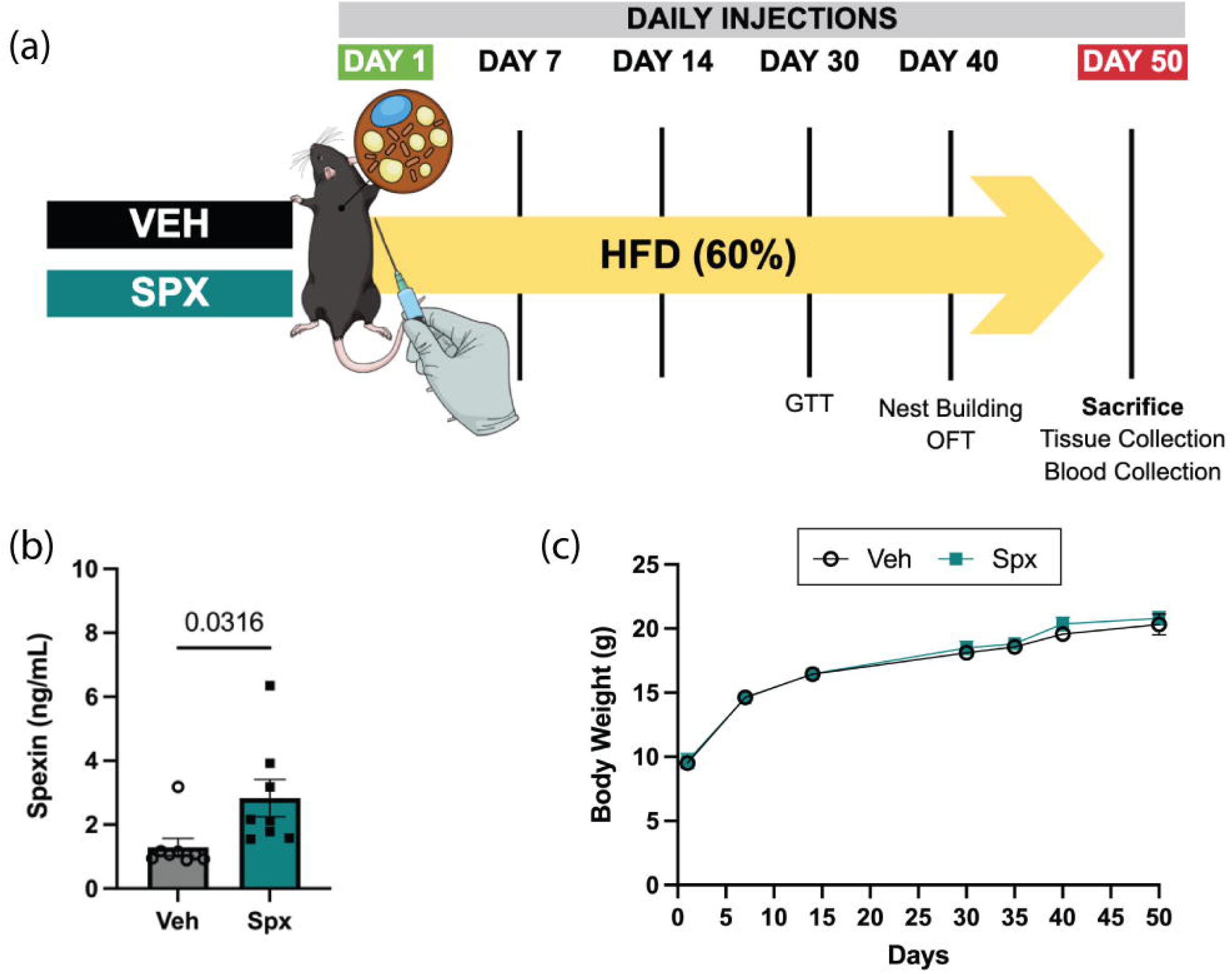

### Spexin improves nest building complexity at room temperature

Nesting behavior in our model of hyperspexinemia was explored after treated animals were observed hoarding paper enrichment material in their cages. Hoarding behaviors in mice are associated with a need to protect resources (food) and conserve body temperature (thermoregulation) [41, 42].

To test the possibility that spexin-treated animals may hoard nesting material for conservation of body temperature, we provided the animals with daily increasing amounts of material. Every interaction with material offered a novel stimulus while nest location preference was scored from 0-5 depending on the location of the nest within the cage (Fig 2a). Nest location has been associated with both direction of air flow within the cage and anxiety [43, 44].

**Fig 2.**
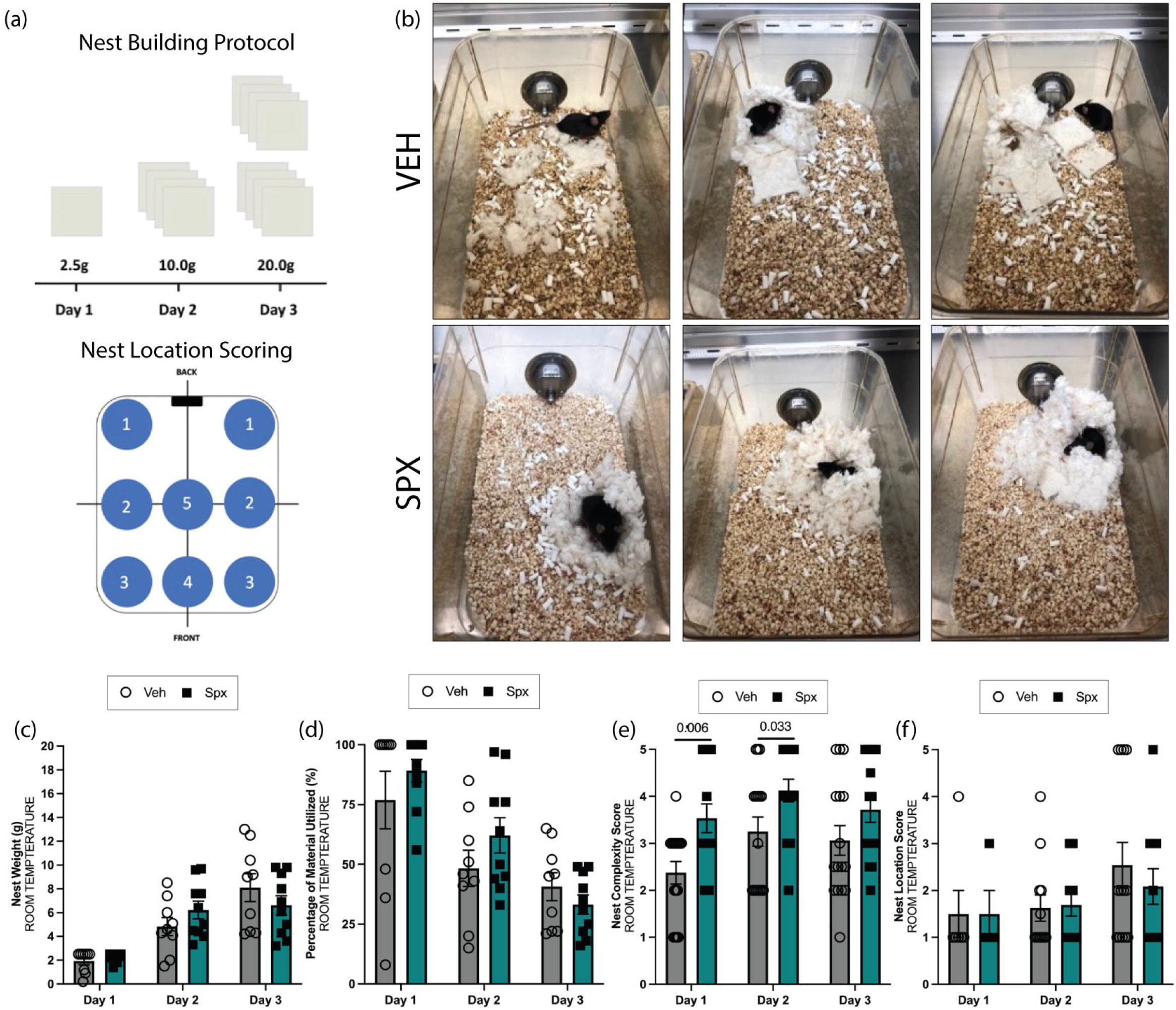

Representative nests were photographed for VEH and SPX after each day of testing at standard room temperatures of 20°C (Fig. 2b). Nest weight was unchanged between either group (Fig. 2c). We then measured the percentage of manipulated material used to build each nest and again found no differences (Fig. 2d). Next, we assessed the complexity of the nests built based upon previously published scoring systems [45, 46, 41]. From Day 1 of testing at the lowest mass of 2.5g of Nestlet™ material to Day 2 of testing with 10.0g of material, spexin-treated animals consistently built more complex nests at standard room temperature with more platform-like, cup shaped, and dome-like nests (Fig. 2e). Although the nests of spexin-treated animals were more complex, the nest location within the cage was not different between the groups, indicating no preferred nest building site at standard room temperature (Fig. 2f).

### Spexin treatment promotes spontaneous exploration of an open field

Frequent interaction with material and repetitive actions such as nest construction are behaviors used to screen for anxiety, compulsive behaviors, and autism-like behaviors in rodents [47, 48]. Note that C57BL/6 animals are inherently anxious compared to other mouse strains [49]. We followed up on our findings using a behavioral assay in a subset of the mice to determine if the improved nest building was attributed to anxiety. We also tested the possibility that spexin-treated animals manipulated nesting material and built nests because spexin treatment increases locomotor activity.

An open field test is a behavioral assay that measures both locomotion and anxiety. In this assay, an individual mouse is placed in the center square of the open field, a situation inherently threatening to a prey species, and its behavior is analyzed for five minutes. Spexin-treatment increased animal reentry into the center square and time spent moving within the center of the field (Fig. 3a,b). Rearing movements were decreased (Fig. 3c) while no differences were observed in other anxiety-related behaviors including stretch attends, grooming, or freezing (Fig. 3d-f). Because spexin-treated animals moved about the open field similarly to the controls, we believe that spexin does not appear to function as locomotion stimulant.

**Fig 3.**
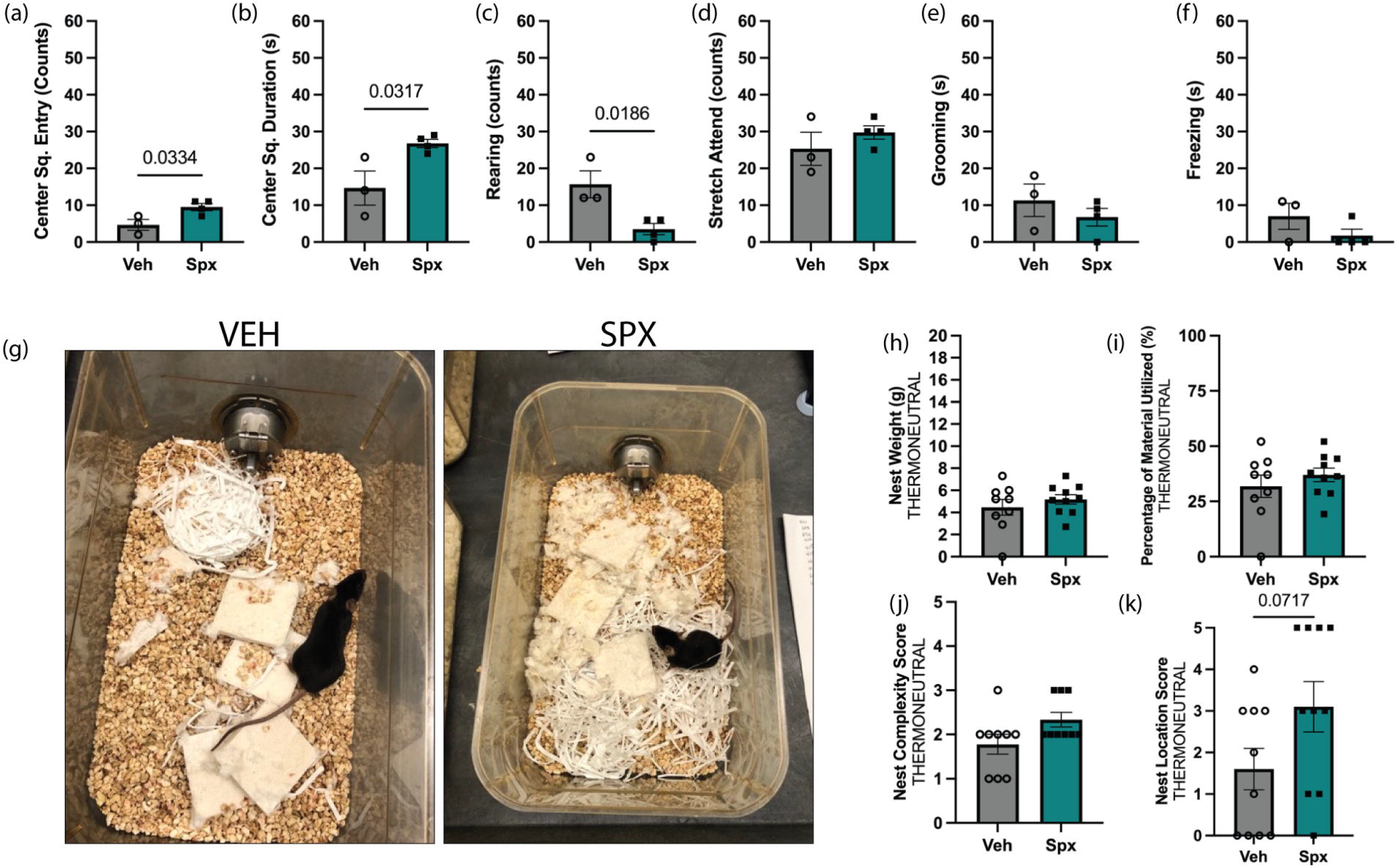

### Thermoneutral temperatures normalize nest building in spexin-treated mice

The animal cages within our animal facility are individually ventilated cages (IVC) where maximal airflow is from the top of the cage. Although IVCs are beneficial for improved airflow, this form of housing at room temperature increases cold stress on animals by increasing activation of brown adipose tissue vacuole size, non-shivering thermogenesis, and sympathetic tone [50, 51]. Based on these conditions, we questioned whether increased nest building complexity was motivated by the need to maintain body temperature. We repeated nest construction assessments at thermoneutral temperatures between 28-30°C[52]. Eighty (80) minutes following their daily spexin injections, animals were exposed to material that consisted of 10.0g of cotton Nestlet™ and 4.0g of paper nesting material. It is worth noting that the shredded paper was novel material for our animals but conducive to nest building by mice [53].

After 16 hours of exposure to thermoneutral temperature (Fig. 3g) and 14g of total nesting material, there were no differences in the amount of material utilized, the percentage of material manipulated to build them, or the complexity of the nests (Fig. 3h-j). Spexin-treated animals chose nest locations that were near the center or far anterior wall (front of cage) instead of the posterior wall or corner of posterior wall (Fig. 4j). These results suggested that the primary motivation for spexin-treated mice to build complex nests was body heat related.

**Fig 4.**
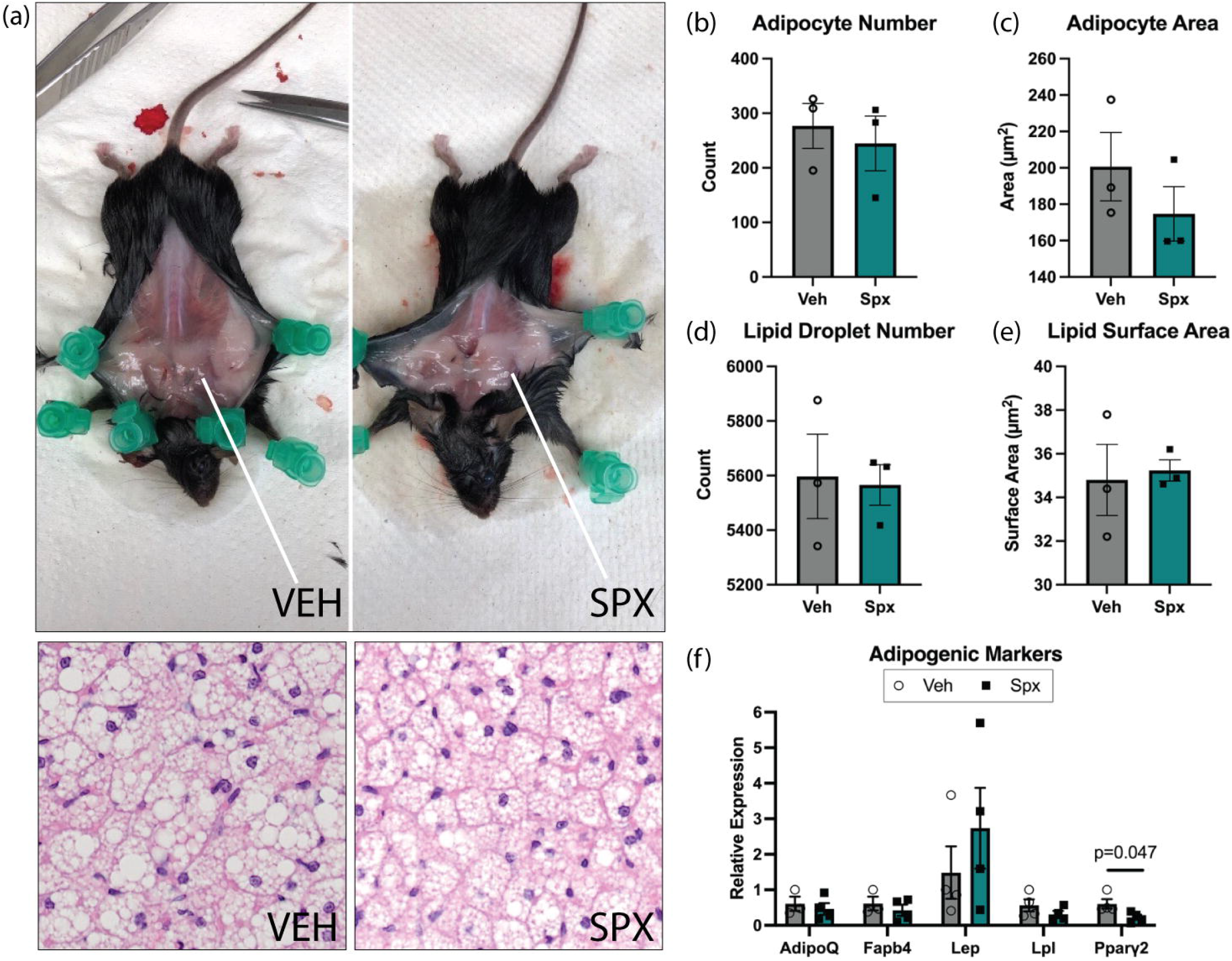

### Spexin reduces thermogenic molecular profile of brown and white adipose tissue

Thermogenesis requires sympathetic innervation of iBAT [54, 55], which in turn triggers browning of other adipose tissue depots such as gonadal white adipose tissue [12, 13]. Given the above results, we investigated if spexin treatment regulates the thermogenic profile of brown adipose tissue. During dissections, we observed that spexin-treated animals appeared to have larger subcutaneous adipose tissue spread across their shoulder blades (Fig 4a). Histological examination of the iBAT however (Fig. 4b-e) revealed that adipocyte number, area, and lipid droplets were not increased and the molecular profile for adipogenic markers was unaffected exclusive of *Pparγ2* mRNA expression that was significantly reduced (p=0.047) in the iBAT of spexin-treated animals (Fig 4f). Upon examination of thermogenic molecular markers in iBAT, our results indicate that treatment downregulated mRNA expression of *Ucp1* while other thermogenic markers such as *Prdm16* (p=0.059), *Dio2* (p=0.076), and *Pgc1α* (p=0.063) trended lower (Fig 5a). Adrenergic receptor activation is a significant pathway in adaptive thermogenesis of brown adipose tissue. It is important for adipocyte differentiation, glucose uptake, lipolysis, and the physiological response to thermal stress to trigger thermogenesis [56]. We assessed adrenoreceptors mRNA expression in brown adipose tissue specifically and found that spexin-treatment did not influence alpha- or beta-adrenoreceptor transcription in BAT (Fig 5b,c). To understand if treatment presented off-target effects from BAT we examined the molecular profile of adrenoreceptors within gonadal white adipose tissue and found that treatment did not affect alpha adrenoreceptors (Fig 5d). Additionally, beta adrenoreceptor type 1 or type 2 mRNA expression was unaffected by treatment except *Adrb3* (p=0.044) also known as β3-adrenergic receptor (β3AR) whose expression was significantly downregulated. Representative images of gWAT appeared to also have less immunodetection of β3AR (Fig. 5f).

**Fig 5.**
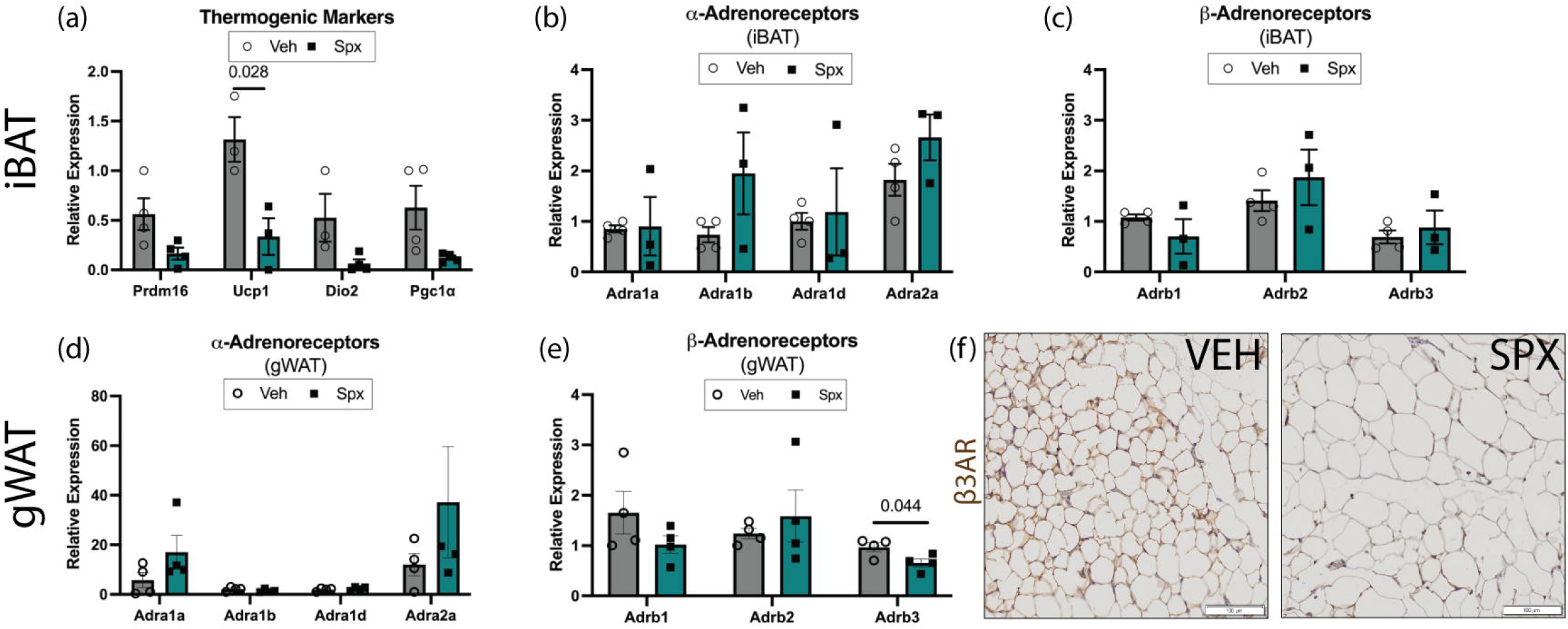

## DISCUSSION

Our study is the first to investigate spexin-treatment as a modulator of thermoregulatory behaviors. Previous spexin studies examined its potential as a treatment for diet-induced obesity. As a biomarker, spexin was the downregulated gene in morbidly obese patients [30]. Within our study, we focused on female animals. Although we were unable to recapitulate the benefits of long-term spexin treatment observed in other studies [30, 31, 57], our study possesses noted differences in both the timing of exposure to high-fat diet, spexin-treatment intervention, and the administration route. Our study began intervention at weaning age when animals were challenged with a high fat diet (HFD). Historically, female C57BL/6 are resistant to diet-induced obesity until after eight weeks of HFD [58, 59].

Behavioral spexin studies have also shown that treatment with the peptide hormone can suppress anxiety-like behaviors across species [32, 33]. Within our cohort, we observed spexin-treated animals hoarding paper material within their enclosures encouraging us to further investigate. We challenged the mice with nest building, an innate behavior that would allow for assessment of motivation and nest structure complexity. These parameters are assessed in animal models for obsessive compulsive-like behaviors [60], anxiety-like behaviors, motor function assessment, thermogenesis [61, 62], and overall wellness [46, 1]. The novelty of our assay was exposure to increasing amounts of material and assessment at both room and thermoneutral temperatures. Room temperature conditions pose thermal stress on animals that prefer warmer temperatures [63]. We found that although spexin-treatment did not increase compulsion to build the treated-animals instead built strikingly complex nests and continued to even when the available material was in excess. When we repeated our nesting protocol at thermoneutral temperatures, we observed normalization of nest-building behavior which confirmed that the behaviors were temperature-driven. To rule out nest building that may manifest as anxiety-like behaviors, animals were placed in the open field behavioral assay. To the contrary, we confirmed the potential for spexin to function as an anxiolytic as others have found [32, 34].

Our findings exploring the molecular and histological profile of adipose tissue and behaviors utilized by mice to conserve body heat suggest a role for spexin in adaptive thermogenesis. Notably, *Ucp1* expression that was downregulated in BAT; *Ucp1* serves as a marker for thermogenic capacity of brown adipose tissue and its expression is enriched in the tissue. When *Ucp1* expression is downregulated or deficient, animals become cold sensitive [64, 65]. Downregulation of β3-adrenergic receptor in WAT may indicate that spexin disrupts the browning potential of WAT and its ability to promote weight loss through WAT depot lipolysis [66, 67].

Overall, this study reveals novel metabolic and behavioral findings that open the door to further investigation of the role of spexin in thermogenesis and thermoregulatory behaviors.

## MATERIALS AND METHODS

### Animals

Pregnant C57BL/6 dams were purchased from The Jackson Laboratories (Bar Harbor, ME) and housed within the Department of Laboratory Animal Resources Facility of the University of Toledo College of Medicine and Life Sciences, Toledo, Ohio, USA under approved IACUC protocol. Animals were given *ad libitum* access to food and water, unless otherwise noted, during metabolic analysis. The dams were on a 12-hr light-dark room schedule. Offspring were weaned at PND 21, and weight-matched into groups for spexin (SPX) or vehicle (Phosphate-buffered saline) treatment. From the date of weaning, animals were placed on a high fat diet with 60% calories from fat (Research Diets, New Brunswick, NJ). Animals were housed four (4) animals per cage with two (2) vehicle animals and two (2) spexin-treated animals.

### Hyperspexinemia Model: Subcutaneous Injection of Spexin

Spexin peptide (Cat. #6090) was purchased from Tocris Bioscience (Bio-Techne Corporation, Minneapolis, MN). To prepare the stock solution, 1.0mg of spexin was dissolved in 1.0 mL of PBS. A final solution of 25 μg/kg was prepared for animal injection studies. Subcutaneous injections began at PND 21. All animals were injected at least 80 minutes prior to lights out in the animal facility (6:00pm). Administration began at PND 21. Animals were subcutaneously injected daily in the scruff between the shoulder blades above the site of interscapular brown adipose tissue (iBAT) with phosphate-buffered solution (PBS, pH=7.4) (VEH or spexin 25 μg/kg dissolved in PBS (SPX). The first day of injections marks Injection Day 1.

### Quantitative Polymerase Chain Reaction

RNA was isolated with a RNeasy Mini Kit (Qiagen, Germantown, Maryland) from dissected tissue and rapidly frozen in liquid nitrogen Quantification with nanodrop. Total RNA (0.75-1.00 μg) was used as template for the High-Capacity cDNA Reverse Transcription Kit (Thermo Fisher) for single-stranded cDNA synthesis. A Bio-Rad Thermal Cycler was used for PCR. Final reaction volumes were diluted 1:5 for qPCR reactions. qPCR reactions were performed as previously described [68]. Primers used for different experiments are listed in Table 1.

### Spexin enzyme immunoassays

Samples of 200μL whole blood were collected by cardiac puncture at the time of sacrifice in microcentrifuge tubes that were allowed to sit at room temperature to coagulate for at least 15 minutes. Samples were centrifuged at 1500 × g for 10 minutes at 4°C. Supernatant (serum) was aliquoted into individually labeled microcentrifuge tubes for a final volume of 50μL and then frozen at 80°C until future analysis. When samples were used for the spexin enzyme immunoassay, serum was thawed to room temperature for 30 minutes, vortexed, and then pipetted in doubles within the EIA kit to measure spexin serum concentration (Phoenix Pharmaceuticals, Burlingame, CA).

### Histology

Hematoxylin and eosin histological staining were performed as previously described [68] to examine morphology of brown adipose tissue in vehicle and spexin-treated animals. Gonadal white adipose tissue samples were embedded in paraffin to preserve then sectioned at a thickness of 5μm. Sections were then deparaffinized and incubated in Tris-EDTA buffer pH 8.0 for 30 minutes at 80ºC. Sections were incubated in blocking buffer containing normal serum of the secondary host, 0.3% hydrogen peroxide, Tris-buffered solution with 0.1% Tween 20, and 1% bovine serum albumin. Tissue sections were then incubated overnight at 4ºC in primary antibody for β-3-adrenergic receptor (Abcam) at a 1:1000 dilution prepared with the same blocking buffer. Following a wash in TBS-T with gentle agitation, samples were then incubated in secondary antibody at a 1:500 dilution and the same blocking buffer solution.

### Automated Image Analysis

Adipocyte area, lipid droplet number, volume, and surface area were assessed with automated image analysis software plugins for Adiposoft (CIMA, University of Navarra) and ImageJ/Fiji Lipid Droplet Counter (Samuel Moll).

### Monitored Nest Building

Nest building behavior was monitored using the Vivarium Animal Intelligence Animal Chamber (Centre Scientific, College Park, Maryland). Media collected of representative nests were assessed blindly by two individuals.

### Nest Complexity Scoring

Nest complexity was scored in a blinded manner by two individuals based on previously described methods in the literature [69, 46, 70, 53, 71]. Complexity was scored from zero (0) to five (5) and based on the percentage of material that was manipulated and the construction style of the nest [69, 46, 70, 72]. Briefly, score = 0, no manipulation of material; Score = 1 ( ≤ 80% manipulation of material); Score = 2 (~80% manipulation of material and placement of material into one area); Score = 3, material placed into an area and hollowed for nestling, Score = 4, flattened nest, hollowed, and higher walls than mouse height with between 75-80% manipulation of material and; Score = 5, ≥80% manipulation of standard cotton Nestlet™ (Ancare Corp., Bellmore, NY) material, dome like structure with entrance and exit, hollowed structure with higher walls.

### Nest Location Preference Scoring

Nest location preference scoring was determined as a score from 0-5, depending on nest position within the cage: Location 1 was designated as either side of the posterior wall of the cage; Location 2 represented the middle of the cage but adjacent to the lateral cage wall; Location 3 represented the front of the cage and greatest distance away from the water source in posterior end of cage; Location 4 represented the center of the anterior wall of the cage; Location 5 represented the center of the cage but not adjacent to a lateral wall. Nest location preference was scored in a blinded manner by two individuals.

### Open Field Test

Anxiety was assessed using the open field test, a five-minute behavioral assay that measures the following parameters: center square reentry, ambulatory activity, defecation, urination, stretch attend postures, rearing, grooming [73–75]. Briefly, a plastic grid arena of 70 cm × 70 cm × 30 cm was cleaned with a 10% bleach solution. After a 60-minute room acclimation period, an individual mouse was placed in the center square of the open field and recorded with a video camera placed directly above the field for five minutes. After the completion of the assay, the open field was cleaned with 10% bleach and air-dried for five minutes. All recorded videos were analyzed for the behavioral parameters by two blinded individuals.

### Statistical Analysis

All graphs were created with GraphPad Prism 8 (GraphPad Software, San Diego, CA). Data is reported as the mean and standard error of the mean. Data were analyzed using student’s t-test for statistical significance and reported as follows: p ≤ 0.05 (*), ≤ 0.01 (**), and ≤ 0.001 (***).

