## Supplementary figures and images for "Spexin modulates molecular thermogenic profile of adipose tissue and thermoregulatory behaviors"

### Supplemental Figure 1

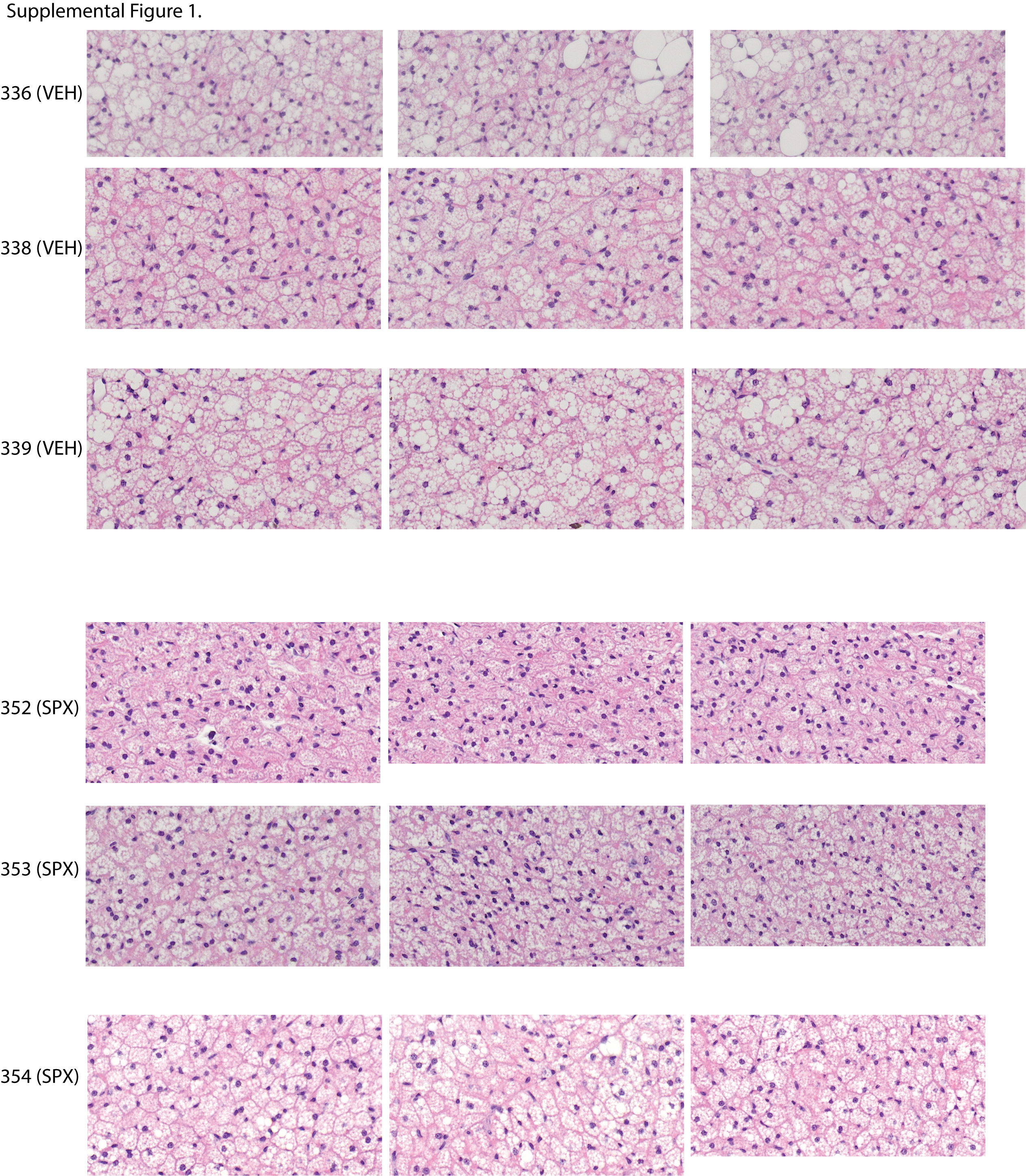
